# Technical Report on Ancient DNA analysis of 27 African Americans from Catoctin Furnace, Maryland

**DOI:** 10.1101/2022.06.12.495320

**Authors:** Éadaoin Harney, Iñigo Olalde, Kari Bruwelheide, Kathryn G. Barca, Roslyn Curry, Elizabeth Comer, Nadin Rohland, Douglas Owsley, David Reich

## Abstract

Catoctin Furnace is an industrial site that functioned from 1776–1903 and was operated at least partially by an enslaved workforce until about 1850, when it shifted to paid European laborers. A cemetery including 35 graves from which the remains of 32 individuals identified with African ancestry were excavated in 1979-1980 during construction of a highway that impacted the site. Since then, the human remains have been under the stewardship of the Smithsonian’s National Museum of Natural History (NMNH). The goal of this report is to document the successful generation of ancient DNA from 27 individuals from the cemetery. The analysis of the Catoctin individuals is part of a long-term study dedicated to restoring knowledge of the identity, origins, and legacy of the enslaved and free African Americans who labored and were buried at Catoctin Furnace^1^.

## Ethics Statement

This analysis was authorized by the NMNH Department of Anthropology Collections Advisory Committee. All work was done in consultation with stakeholders from the Catoctin Furnace Historical Society (CFHS) and the African American Resources, Cultural and Heritage Society (AARCH) in Frederick, Maryland. During the course of this work, members of CFHS successfully traced the lineages of two African American individuals who labored at Catoctin to some of their living descendants. Any further studies using the genetic data included in this report should consider consultation with stakeholders from CFHS, AARCH, and direct descendant families.

## Methods

We sampled the Catoctin skeletal remains in two batches following standard approaches (Table 1). Briefly:

**Table 1.** Results of successful genetic screening of 27 Catoctin individuals.

| Burial ID | Harvard Laboratory ID | Smithsonian ID | Sampling Batch | Percent Endogenous | mtDNA coverage | Rate of mtDNA match to consensus sequence | mtDNA haplogroup | Coverage across all 1240k targets | Number of SNPs hit on autosomal 1240k targets | Damage rate in first nucleotide on sequences overlapping 1240k targets | Sex ratio [Y/(Y+X) counts] | Genetic Sex | Y Haplogroup | X chromosome contamination estimate (for males with >200 reads aligning to X chromosome targets) | Family | Number of ref (T): alt (A) alleles observed at SNP rs334 | Notes |
| --- | --- | --- | --- | --- | --- | --- | --- | --- | --- | --- | --- | --- | --- | --- | --- | --- | --- |
| 1 | I15330 | 387788 | 2 | 2.4% | 152.50 | 0.994 ± 0.006 | L3e2a1b1 | 0.72 | 510,487 | 0.100 | 0.401 | M | E1b1a1a1a1c1b | 0.005±0.006 | A | 5:0 |  |
| 2 | I15331 | 387789 | 2 | 13.9% | 532.40 | 0.977 ± 0.010 | L3e2a1b1 | 2.55 | 774,043 | 0.063 | 0.408 | M | E1b1a1a1a1c1b1 | 0.005±0.003 | A | 22:0 |  |
| 3 | I15332 | 387790 | 2 | 23.1% | 93.17 | 0.990 ± 0.007 | L3e2a1b1 | 0.45 | 383,530 | 0.090 | 0.015 | F | .. | .. | A | 0:0 |  |
| 4 | I8084 | 387791 | 1 | 3.5% | 33.50 | 0.964 ± 0.009 | L0a1b1a | 0.28 | 261,081 | 0.044 | 0.446 | M | A1b1 | - |  | 0:0 |  |
| 5 | I8085 | 387792 | 1 | 10.4% | 102.00 | 0.993 ± 0.004 | L3e1 | 1.07 | 515,648 | 0.023 | 0.435 | M | E1b1a1a1 | 0.005±0.005 | D | 4:0 |  |
| 6 | I8086 | 387793 | 1 | 5.5% | 43.90 | 0.991 ± 0.006 | L2a1a1 | 1.04 | 510,367 | 0.035 | 0.438 | M | E1b1a1a1a1c1b2a | 0.002±0.003 |  | 6:0 |  |
| 7 | I15333 | 387794 | 2 | 4.6% | 110.20 | 0.995 ± 0.004 | L3e1a1a | 1.08 | 594,078 | 0.117 | 0.01 | F | .. | .. |  | 3:0 |  |
| 8 | I8087 | 387795 | 1 | 11.3% | 76.70 | 0.989 ± 0.006 | L3e2a1b1 | 1.29 | 567,644 | 0.038 | 0.429 | M | E1b1a1a1a1a | 0.008±0.005 |  | 8:0 |  |
| 9 | I15334 | 387796 | 2 | 7.0% | 270.60 | 0.989 ± 0.007 | L3f1b1a | 1.72 | 709,865 | 0.071 | 0.011 | F | .. | .. |  | 4:0 |  |
| 10 | I8088 | 387797 | 1 | 8.0% | 83.00 | 0.992 ± 0.004 | L3f1b3 | 1.14 | 529,609 | 0.030 | 0.438 | M | R1b1a1b1a1a2c1a1h2a~ | 0.005±0.004 |  | 11:0 |  |
| 11 | I8089 | 387798 | 1 | 23.5% | 113.00 | 0.996 ± 0.002 | L2b1a3 | 1.11 | 505,029 | 0.054 | 0.019 | F | .. | .. | B | 5:0 |  |
| 12 | I15335 | 387799 | 2 | 3.2% | 161.60 | 0.983 ± 0.006 | L2b1a3 | 0.50 | 406,464 | 0.072 | 0.012 | F | .. | .. | B | 1:0 |  |
| 13 | I8090 | 387800 | 1 | 5.4% | 1.98 | - | L2b1a3 | 0.02 | 24,971 | 0.031 | - | M | E2b | - | B | 4:0 | Damage Restricted |
| 14 | I8091 | 387801 | 1 | 8.8% | 87.10 | 0.957 ± 0.008 | L4b2b1 | 0.59 | 376,829 | 0.028 | 0.026 | F | .. | .. |  | 1:0 |  |
| 15 | I8092 | 387802 | 1 | 43.3% | 286.00 | 0.999 ± 0.001 | L2a1+143+16189 (16192)+@16309 | 1.88 | 640,373 | 0.035 | 0.423 | M | E1b1a1a1a2a1a | 0.008±0.004 | C | 12:0 |  |
| 17 | I15336 | 387804 | 2 | 3.8% | 97.29 | 0.991 ± 0.007 | L3e2 | 0.64 | 494,269 | 0.043 | 0.389 | M | E1b1a1a1a1c1a1 | 0.026±0.012 | E | 0:1 |  |
| 18 | I8093 | 387805 | 1 | 12.3% | 108.00 | 0.997 ± 0.002 | L2a1+143+16189 (16192)+@16309 | 0.95 | 508,736 | 0.025 | 0.019 | F | .. | .. | C | 4:0 |  |
| 19 | I15337 | 387806 | 2 | 3.0% | 72.80 | 0.980 ± 0.011 | L3e2 | 0.52 | 422,520 | 0.057 | 0.011 | F | .. | .. | E | 0:3 |  |
| 22 | I8094 | 387809 | 1 | - | 66.80 | 0.989 ± 0.006 | L3d1b3 | 1.63 | 694,538 | 0.080 | 0.402 | M | E1b1a1a1a1c1a1a3a1d1 | 0.010±0.004 |  | 0:5 |  |
| 23 | I15339 | 387810 | 2 | 30.5% | 18.81 | 0.995 ± 0.005 | L2a1+143+16189 (16192)+@16309 | 0.08 | 86,568 | 0.056 | 0.014 | F | .. | .. | C | 0:0 |  |
| 24 | I15340 | 387811 | 2 | 11.6% | 195.20 | 0.996 ± 0.005 | L3e2a1b1 | 0.98 | 612,083 | 0.081 | 0.402 | M | E1b1a1a1a1c1a1 | 0.016±0.007 | A | 3:0 |  |
| 26 | I15341 | 387813 | 2 | 0.4% | 1.54 | - | L2c | 0.01 | 11,874 | 0.126 | - | M | E1b1a1~ | - |  | 0:0 | Damage Restricted |
| 28 | I15338 | 387814 | 2 | 1.1% | 17.57 | 0.989 ± 0.008 | L2b1a3 | 0.03 | 39,262 | 0.119 | 0.018 | F | .. | .. |  | 0:0 |  |
| 32 | I15342 | 387816 | 2 | 9.0% | 39.52 | 0.991 ± 0.007 | J1b1a1a | 0.31 | 300,899 | 0.126 | 0.395 | M | R1a1a1 | 0.002±0.007 |  | 0:0 |  |
| 33 | I8095 | 387817 | 1 | - | 128.00 | 0.997 ± 0.002 | L2a1+143+@16309 | 3.12 | 774,454 | 0.071 | 0.408 | M | E1b1a1a1a1c2c | 0.005±0.002 |  | 19:0 |  |
| 34 | I8096 | 387818 | 1 | - | 102.00 | 0.991 ± 0.006 | L3e1 | 2.07 | 739,557 | 0.037 | 0.402 | M | R1b1a1b1a1a2c1 | 0.003±0.002 | D | 10:0 |  |
| 35 | I8097 | 387819 | 1 | - | 151.00 | 0.994 ± 0.004 | L3e1 | 3.87 | 787,170 | 0.062 | 0.008 | F | .. | .. | D | 33:0 |  |

Dashes (-) indicate cases where too few aligned reads were generated to calculate the metric. Two dots (..) indicate that the metric was not calculated, as the individual is female.

1. For each individual, we sampled powder from the petrous portion of the temporal bone, using a sandblaster and mixer mill when sampling from disarticulated temporals^3,4^ (batch 1) and a cranial based drilling approach for intact skulls^2^ (batch 2).
2. We extracted DNA from approximately 37 mg of bone powder for each petrous bone sample^5^.
3. We converted each DNA extract into a single barcoded, partially UDG-treated Illumina sequencing library^6^.
4. We increased the proportion of human DNA in the libraries through targeted enrichment capture using oligonucleotide probes aligning to the mitochondrial genome and 1.2 million SNPs in the nuclear genome^7– 9^, which we refer to as the “1240k targets”. For libraries in batch 1, the mitochondrial and 1240k capture was performed separately. For batch 2, the probes were pooled into a single capture reagent.
5. We sequenced the enriched libraries (and performed a small amount of shotgun sequencing of the unenriched libraries) using either an Illumina HiSeq10 or NextSeq500 instrument, with 2×101 or 2×76 cycles, respectively, and an additional 2×7 cycles to sequence the library indices.
6. We performed bioinformatic processing of the sequenced DNA using a custom software (https://github.com/DReichLab/ADNA-Tools), aligning the sequences to the mitochondrial consensus sequence (RSRS)^10^ and human reference genome (version hg19).

These procedures are like those used to generate similar data for other Reich laboratory papers.

## Results

### (1) DNA Authenticity

We successfully obtained DNA from all 27 of the sampled Catoctin individuals. Metrics describing the amount or quality of DNA obtained from each individual are reported in Table 1. Throughout the report, we use the following convention to refer to each of the Catoctin individuals: “Burial ID – Harvard Laboratory ID,” for example, 1-I15330.

To assess DNA authenticity, we considered 3 standard metrics:

a) <u>*C-to-T substitution rate*</u>: All Catoctin individuals exhibited a C-to-T substitution rate of at least 2.3% at the terminal end of sequenced reads. Typically, a C-to-T substitution rate of 3% at the terminal end of molecules is used as a standard minimum threshold when assessing the authenticity of ancient DNA molecules. However, this threshold was defined based on DNA sampled from more ancient individuals (typically thousands of years old). Damage in ancient DNA accumulates over time. Based on the relatively recent age of the Catoctin individuals, we considered all individuals to have sufficient evidence of C-to-T substitution to be suitable for further analysis.

b) <u>*Mitochondrial contamination*</u>: All Catoctin individuals exhibited a mitochondrial match to consensus rate that exceeds the standard minimum threshold of 95%, as determined by contamix v1.0-12^11^.

c) <u>*X-chromosome contamination*</u>: We determined the X-chromosome contamination rate for all individuals that we identified as genetically male and that had over 200 sequences that aligned to targeted single nucleotide polymorphisms (SNPs) on the X-chromosome. A minimum contamination rate of below 3% is typically used to determine DNA authenticity using the tool ANGSD, and all the Catoctin individuals for whom this metric could be calculated received contamination estimates below this threshold.

Although all 27 sequenced libraries passed these screening metrics, the lower bound of the 95% mitochondrial match to consensus rate and the upper bound of the X-chromosome contamination rate fell outside of the acceptable limits for two libraries (13-I8090 and 24-I15340). Out of an abundance of caution, we restricted all analyses of these individuals (including the metrics reported in Table 1) to sequences that show evidence of having C-to-T damage, as determined by the tool PMDtools^12^.

### (2) Determination of Genetic Sex, Mitochondrial and Y-chromosome haplogroups

We inferred the genetic sex of each Catoctin individual by considering the observed ratio of sequences that align to the X and Y chromosomes out of the shotgun sequenced reads^13^. We identified 11 genetically female and 16 genetically male individuals (Table 1). We inferred the mitochondrial DNA haplogroup of each individual using haplogroup2, with Phylotree version 17^14^. We inferred a Y haplogroup by identifying the most derived mutation, using nomenclature defined by the International Society of Genetic Genealogy (ISOGG) version 14.76 (April 2019).

Most of the Catoctin individuals have mitochondrial DNA and Y chromosome haplogroups that are at highest frequency among populations with African ancestry. Three individuals (10-I8088, 32-I15342, and 34-I18096) were assigned Y haplogroups that are at high frequency among European populations (falling within subclades of the R1a and R1b lineages), while only a single individual (32-I15342) was assigned a European-associated mitochondrial haplogroup. The observation of a greater number of European-associated Y haplogroups than mitochondrial haplogroups mirrors known patterns of sex-biased admixture among African Americans, much of which is the result of rape or cajoled sexual relations between white men and enslaved Black women^15,16^.

### (3) Identification of genetic relatives

We used a previously described approach^17^ to identify genetic relatives of 1^st^–3^rd^ degree. The method measures the rate at which two randomly chosen sequences from two individuals differ from each other when they overlap at the same position and compares it to the rate at which two randomly chosen sequences from the same population differ. Under the assumption that a person’s two parents are not closely related, the rate of differences between their two chromosomes can be used to establish a baseline expectation for the difference rate between two unrelated individuals in a population. A 1^st^-degree relative is expected to possess half of that difference rate, a 2^nd^-degree relative is expected to have a quarter of that difference rate, and a 3^rd^-degree relative is expected to have an eighth of that difference rate. We identified 5 genetic family groups:

a) Family A has 4 members. Individual 3-I15332 is the mother of two sons, 1-I15330 and 2-I15331. All three individuals have a 2^nd^–3^rd^-degree relationship with individual 24-I15340.

b) Family B has 3 members. Individual 11-I18089 has a 2^nd^–3^rd^-degree relationship with individuals 13-I8090 and 12-I15335, who are genetically unrelated to one another. Individual 28-I15338 may also be related to members of Family B, but we do not have sufficient coverage to make a definitive determination.

c) Family C has 3 members. Individuals 15-I8092 and 18-I8093 are siblings. Both individuals also have a 1^st^-degree relationship with individual 23-I15339, but we could not determine whether this individual is their mother or sister.

d) Family D has 3 members. Individual 35-I8097 is the mother of individual 34-I8096 and the sister of individual 5-I8085.

e) Family E has 2 members, siblings 17-I15336 and 19-I15337.

### (4) Ancestry modeling

We used the tool qpAdm^18^ (version 960) to estimate the proportion of African, European, and Indigenous American ancestry of each of the Catoctin individuals. We selected the source populations YRI.SG (Yoruba in Ibadan, Nigeria), GBR.SG (British in England and Scotland), and Pima.SDG (Indigenous population from northwestern Mexico) to represent each of these broadly defined ancestries, respectively. Since the goal of this analysis was to generate broad estimates of each of these continental level ancestries, we chose Mbuti.SDG (Indigenous population from the Congo), Khomani_San.DG (Aboriginal population from South Africa), CHB.SG (Han Chinese in Beijing, China), and FIN.SG (Finnish in Finland) to serve as reference populations, as we believe that they are unlikely to be more closely related to the Catoctin individuals than the selected source populations. This qpAdm model is therefore likely to be highly permissive with respect to the true sources of African, European, and Indigenous American ancestry observed among the Catoctin individuals. The results of this analysis are reported in Table 2. We considered all models with p-values >0.01 and estimated ancestry proportions that fall within 3-standard errors of the range 0–1 to be plausible.

**Table 2.** qpAdm estimates of African, European, and Indigenous American-related ancestry.

| Burial ID | Harvard Laboratory ID | Smithsonian ID | p-value | Proportion of Ancestry Assigned to YRI.SG | Proportion of Ancestry Assigned to GBR.SG | Proportion of Ancestry Assigned to Pima.SDG | Standard Error YRI.SG | Standard Error GBR.SG | Standard Error Pima.SG | Notes |
| --- | --- | --- | --- | --- | --- | --- | --- | --- | --- | --- |
| 1 | I15330 | 387788 | 0.092 | 0.879 | 0.111 | 0.011 | 0.011 | 0.013 | 0.011 | Pass |
| 2 | I15331 | 387789 | 0.054 | 0.882 | 0.092 | 0.026 | 0.010 | 0.012 | 0.011 | Pass |
| 3 | I15332 | 387790 | 0.063 | 0.890 | 0.135 | -0.025 | 0.011 | 0.013 | 0.010 | Pass |
| 4 | I8084 | 387791 | 0.054 | 0.983 | 0.026 | -0.009 | 0.007 | 0.011 | 0.011 | Pass |
| 5 | I8085 | 387792 | 0.096 | 0.937 | 0.057 | 0.006 | 0.008 | 0.010 | 0.009 | Pass |
| 6 | I8086 | 387793 | 0.014 | 1.015 | 0.011 | -0.026 | 0.006 | 0.009 | 0.009 | Pass |
| 7 | I15333 | 387794 | 0.026 | 1.020 | -0.017 | -0.004 | 0.006 | 0.008 | 0.009 | Fail: YRI.SG ancestry > 1+ 3xSE |
| 8 | I8087 | 387795 | 0.030 | 1.011 | -0.005 | -0.006 | 0.006 | 0.009 | 0.009 | Pass |
| 9 | I15334 | 387796 | 0.045 | 1.004 | -0.007 | 0.003 | 0.006 | 0.009 | 0.009 | Pass |
| 10 | I8088 | 387797 | 0.370 | 0.558 | 0.445 | -0.003 | 0.015 | 0.017 | 0.012 | Pass |
| 11 | I8089 | 387798 | 0.036 | 1.013 | -0.008 | -0.005 | 0.006 | 0.008 | 0.008 | Pass |
| 12 | I15335 | 387799 | 0.019 | 1.013 | -0.001 | -0.012 | 0.006 | 0.008 | 0.009 | Pass |
| 13 | I8090 | 387800 | 0.144 | 0.935 | -0.005 | 0.070 | 0.019 | 0.029 | 0.030 | Pass |
| 14 | I8091 | 387801 | 0.067 | 0.770 | 0.251 | -0.021 | 0.011 | 0.014 | 0.010 | Pass |
| 15 | I8092 | 387802 | 0.036 | 0.943 | 0.078 | -0.021 | 0.010 | 0.012 | 0.010 | Pass |
| 17 | I15336 | 387804 | 0.026 | 0.957 | 0.051 | -0.008 | 0.008 | 0.010 | 0.009 | Pass |
| 18 | I8093 | 387805 | 0.009 | 0.974 | 0.056 | -0.029 | 0.010 | 0.011 | 0.009 | Fail: p < 0.01 |
| 19 | I15337 | 387806 | 0.009 | 0.946 | 0.061 | -0.008 | 0.009 | 0.011 | 0.010 | Fail: p < 0.01 |
| 22 | I8094 | 387809 | 0.022 | 0.900 | 0.110 | -0.010 | 0.010 | 0.012 | 0.009 | Pass |
| 23 | I15339 | 387810 | 0.002 | 0.984 | 0.043 | -0.028 | 0.013 | 0.018 | 0.017 | Fail: p < 0.01 |
| 24 | I15340 | 387811 | 0.039 | 1.025 | -0.001 | -0.024 | 0.006 | 0.008 | 0.008 | Fail: YRI.SG ancestry > 1 + 3xSE |
| 26 | I15341 | 387813 | 0.362 | 0.808 | 0.215 | -0.024 | 0.029 | 0.046 | 0.046 | Pass |
| 28 | I15338 | 387814 | 0.242 | 0.951 | 0.040 | 0.010 | 0.015 | 0.023 | 0.022 | Pass |
| 32 | I15342 | 387816 | 0.468 | 0.436 | 0.587 | -0.023 | 0.010 | 0.014 | 0.012 | Pass |
| 33 | I8095 | 387817 | 0.022 | 0.984 | 0.009 | 0.007 | 0.006 | 0.008 | 0.008 | Pass |
| 34 | I8096 | 387818 | 0.015 | 0.886 | 0.109 | 0.006 | 0.009 | 0.011 | 0.009 | Pass |
| 35 | I8097 | 387819 | 0.042 | 0.962 | 0.040 | -0.002 | 0.007 | 0.009 | 0.008 | Pass |

Using this approach, we successfully modeled the ancestry of 22 of the Catoctin individuals—an additional 2 individuals have YRI.SG-related ancestry in excess of 100% plus 3 standard errors (Table 2), which indicates they likely have full African ancestry, but that this ancestry is not well represented by the source population YRI.SG. Among the 22 individuals whose ancestry we successfully modeled, there is a great deal of heterogeneity among the Catoctin individuals with respect to European ancestry, and 7 of the Catoctin individuals can be modeled as having no European ancestry (i.e., the amount of ancestry assigned to GBR.SG is within 1 standard error of 0). Conversely, 13 of the Catoctin individuals could not be modeled without European-related ancestry (i.e., the amount of ancestry assigned to GBR.SG is greater than 3 times the assigned standard errors). While many of the Catoctin individuals were modeled as having a small proportion of Indigenous American-related ancestry, all estimates were within 3 standard errors of 0, and therefore it could not be confidently determined whether any of the Catoctin individuals have Indigenous American related ancestry via this analysis.

### (5) Limited evidence of possible sickle cell disease or trait

We counted the number of unique alternative and reference alleles that aligned to the *rs334* SNP. Individuals that possess two copies of the alternative allele, A, at this position have the red blood cell disorder, Sickle Cell Disease, while individuals who are heterozygous at this position are considered carriers of Sickle Cell Trait and are less susceptible to malaria^19^. We observed the causal A allele in three of the Catoctin individuals, siblings 17-I15336 and 19-I15337, and individual 22-I8094 (Table 1). No sequences carrying the reference, T, were observed for any of these individuals. However, no more than 5 reads aligned to this position for any of these individuals. Therefore, it cannot be stated with confidence whether they had Sickle Cell Disease or instead were carriers of Sickle Cell Trait.

## Conclusion

These results demonstrate that the Catoctin Furnace African American cemetery served as a burial ground for both related and unrelated African American individuals. These individuals possessed heterogeneous amounts of African and European ancestry, and three individuals may have suffered from Sickle Cell Disease or were carriers of the trait.

These genetic results are featured in the Catoctin Furnace African American Cemetery Interpretive Trail, which was unveiled in 2020, and in an exhibit of forensic facial reconstructions of two individuals (15-I8092 and 35-I8097) that are on display at the Catoctin Furnace Museum of the Ironworker. These facial reconstructions were unveiled June 24^th^, 2021, in Frederick, Maryland, and several co-authors of this report (E.H., K.B., E.C., and D.O.) were present and participated in the presentation of the findings. This report serves as a formal record of the genetic analyses. The Reich laboratory often increases the quality of DNA data from individuals using bone powder and DNA extracts stored in the laboratory after sampling (all bone samples from Catoctin have been returned to the NMNH for long-term stewardship). If additional data are produced in the future allowing higher resolution analyses for the same individuals, the data will be added to the online database (see Data Availability section). The present document serves as the formal reference for those data.

## Data Availability

Unaligned reads in fastq format in addition to autosomal and mtDNA BAMS are available from the European Nucleotide Archive under accession number PRJEB52230. Genotype data are available from the Reich lab datasets webpage (https://reich.hms.harvard.edu/datasets).

## Acknowledgments

We are grateful to Linda Heywood, Henry Louis Gates Jr., Jakob Sedig, Kendra Sirak, and John Thornton for conversations; Rebecca Bernardos, Adam Micco, Matthew Mah, Shop Mallick, and Zhao Zhang for sample handling and bioinformatics support; and Nicole Adamski, Nasreen Broomandkhoshbacht, Matthew Ferry, Lijun Qiu, Kristin Stewardson, Noah Workman, and Fatma Zalzala for wet laboratory work. The ancient DNA research was funded by John Templeton Foundation grant 61220 and the Howard Hughes Medical Institute. In 2016, a $14,000 Maryland Heritage Areas Authority grant was awarded to CFHS to fund forensic research for the African American cemetery

## Notes

### Competing Interest Statement

Eadaoin Harney is an employee of 23andMe. Roslyn Curry was an intern at 23andMe in summer 2021.

